# MDPath: Unraveling Allosteric Communication Paths of Drug Targets through Molecular Dynamics Simulations

**DOI:** 10.1101/2025.03.27.645168

**Authors:** Niklas Piet Doering, Marvin Taterra, Marcel Bermúdez, Gerhard Wolber

## Abstract

Understanding allosteric communication in proteins remains a critical challenge for structure-based, rational drug design. We present *MDPath*, a Python toolkit for analyzing allosteric communication paths in molecular dynamics simulations using NMI-based analysis. We demonstrate *MDPath*’s ability to identify both established and novel GPCR allosteric mechanisms using the β_2_-adrenoceptor, adenosine A_2A_ receptor, and µ-opioid receptor as model systems. The toolkit reveals ligand-specific allosteric effects in β_2_-adrenoceptor and MOR, illustrating how protein-ligand interactions drive conformational changes. Analysis of ABL1 kinase in complex with allosteric and orthosteric inhibitors demonstrates the broader applicability of the approach. Ultimately, *MDPath* provides an open-source framework for mapping allosteric communication within proteins, advancing structure-based drug design (https://github.com/wolberlab/mdpath).

## 1 Introduction

In recent years, significant progress has been made in predicting ligand binding using molecular docking [1], free energy perturbation [2], and in analyzing ligand interactions through static and dynamic 3D pharmacophores [3]. However, identifying the allosteric networks that drive and stabilize conformational changes remains a major challenge in structure-based drug discovery (Fig. 1). Recent approaches, including AI-driven predictions[4] and residue-residue contact scores[5–7], have enhanced our mechanistic understanding. However, most of these methods are primarily effective for isolated switches or smaller systems, limiting their ability to elucidate full patterns in larger systems.

**Fig. 1.**
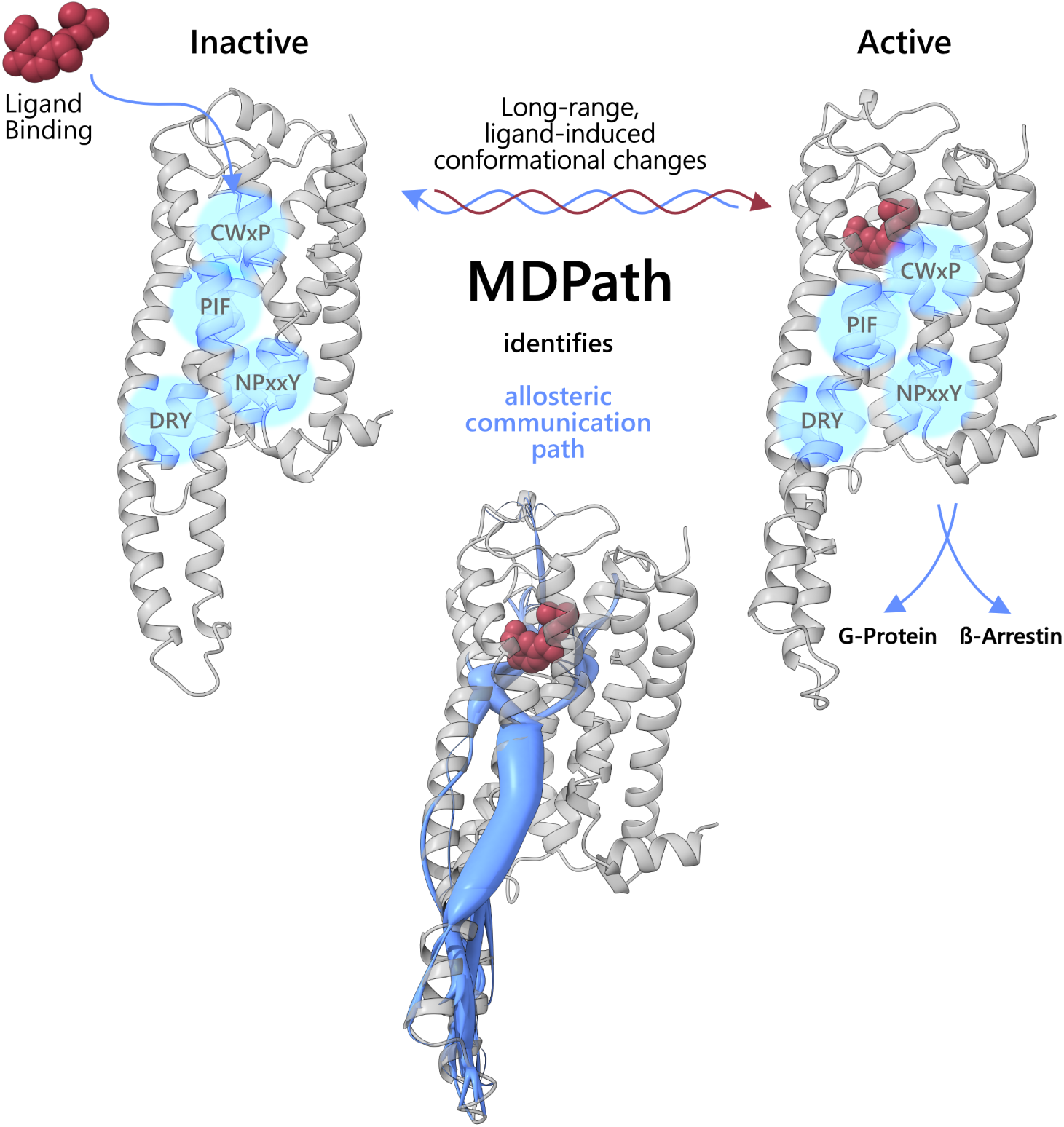
GPCR transition between inactive (left) and active (right) states upon ligand (red) binding, leading to downstream signaling through G proteins or β-arrestins. However, the full molecular mechanism by which conformational changes propagate through the receptor remains unclear. Key conserved motifs (CWxP, PIF, NPxxY, and DRY) involved in GPCR activation are highlighted (blue circles). MDPath, an approach for mapping allosteric communication (middel) through NMI and graph-based path tracking, identifies allosteric communication paths (blue) that connect and stabilize distant protein regions. This allows MDPath to track the effects of drug binding sites on distant effector regions, enabling rationalization beyond protein-ligand interactions.

A promising approach to address this challenge was introduced by McClendon et al. in 2009, which uses mutual information (MI) between residue dihedral angle movements to map these communication networks.[8] Mutual information is a measure for the dependence between two random variables - in this case, the dynamic movements of the dihedral angles of the protein backbone.[9] By measuring the correlation between the motion of one residue and that of another, mutual information provides insight into the potential functional connectivity between distant regions of a protein, even in cases where direct interactions are not possible due to spatial separation. This allows the identification of long-range allosteric paths, capturing a complete network that drives conformational shifts and protein function.[6–8, 10, 11]

Understanding allosteric communication in proteins is crucial for modern drug development, since it requires mechanistic understanding on atomistic level. This is particularly evident in G protein-coupled receptors (GPCRs), which account for 35% of approved drugs.[12] In GPCRs, allosteric communication paths connect ligand-binding sites to intracellular effector domains, facilitating precise signal propagation.[13] These pathways involve conserved microswitches such as the CWxP, PIF, NPxxY, and DRY motifs (Fig. 1), which play a critical role in the process but do not fully define the connection. Instead, a complex network of structural rearrangements extends beyond these motifs, orchestrating the transmission of conformational changes required for receptor activation.[5, 13, 14] The therapeutic significance of GPCRs is demonstrated by well-characterized systems: the β_2_-adrenoceptor, a primary target for asthma.[15]; the A_2A_ adenosine receptor, which has emerged as a targeted for inflammatory conditions and neurological disorders[16]; and the µ-opioid receptor (MOR), where improved understanding of allosteric mechanisms could lead to safer analgesics with reduced adverse effects like tolerance and respiratory depression[17].

Besides GPCRs, there are many other drug targets for which allosteric mechanisms have been reported and provide the possibility to develop specific modulators. One example is BCR-ABL, a constitutively active tyrosine kinase arising from the Philadelphia chromosome translocation[18, 19]. While normal ABL1 is regulated through an N-terminal myristoyl-mediated autoinhibition mechanism, BCR-ABL lacks this regulatory element, leading to uncontrolled kinase activity and leukemia development[20, 21]. The therapeutic success of targeting distinct conformational states is demonstrated by two drug classes: ATP-competitive inhibitors like bosutinib that stabilize an inactive conformation[22], and newer allosteric inhibitors such as asciminib that mimic the natural autoinhibition mechanism by targeting the myristoyl pocket[23, 24].

Recognizing the critical role of allosteric communication, we have developed *MDPath* (Fig. 1), a novel open-source tool that integrates normalized mutual information (NMI)[8] with graph-based path tracking to map, analyze, and visualize allosteric coupling. Beyond identifying global networks, *MDPath* traces specific allosteric paths linked to key residues, enabling direct correlation of ligand binding or mutations with conformational shifts. By providing a systematic framework for characterizing allosteric paths, *MDPath* not only deepens our understanding of known molecular mechanisms but also holds the potential to uncover novel paths underlying previously uncharacterized mechanisms.

## 2 Results

### 2.1 MDPath Implementation

MDPath implements a comprehensive computational workflow for characterizing allosteric communication paths in MD data. The core analysis pipeline integrates NMI calculations for residue correlation with graph-based topology analysis and a novel direct ligand-based path tracking system. The workflow leverages state-of-the-art libraries including MDAnalysis[28], NumPy[29], Pandas[30], SciPy[31], scikit-learn[32], and NetworkX[33] for efficient processing and analysis of MD trajectories.

We employed dihedral angle analysis as the foundation of our computational approach, calculating φ-dihedral angles for all protein residues from the input trajectory data. The φ angle was chosen because it is present in every amino acid, unlike the variable χ angles, and in the context of a protein, φ and ψ angles show correlated behavior. The mutual information between residue movements is computed through histogram-based analysis using *NumPy* [29], treating joint distributions as contingency tables. The mutual information values are normalized using the geometric mean of individual residue entropies calculated via *SciPy* [31], providing robust measures of correlation between residue movements. MDPath constructs a detailed residue interaction network where nodes represent individual residues and edges connect proximate residues (within 5 Å), weighted by their NMI values. The pipeline implements Dijkstra’s algorithm[34] to identify paths between residues separated by at least 12 Å, focusing on paths with maximum cumulative NMI. To increase robustness and reduce noise, we are using a ranking system that selects the top 500 paths based on total NMI scores. Hierarchical clustering is then employed to delineate distinct allosteric communication paths, using heavy atom distances between residues across different paths as a metric for path overlap. The optimal number of clusters is determined via silhouette score analysis, ensuring cluster separation and cohesion.

For visualization, MDPath integrates with *NGLView* [25] for interactive *Jupyter* notebook representation and a *PyMOL*[26] script for structural rendering. Path representations are anchored to backbone α-carbon atoms, with connection radii scaled by path frequency within clusters, providing an intuitive depiction of allosteric network significance. Additionally, the workflow supports an STL export of the paths as precomputed splines for 3D modeling applications such as Blender[35] or visualization in *ChimeraX* [27].

### 2.2 Recognition of Conserved Motifs in GPCRs

To initially validate *MDPath*, we assessed its capacity to identify conserved motifs across class A GPCRs, focusing on the β_2_-adrenoceptor, adenosine A_2A_ receptor, and MOR with agonists, antagonists and inverse agonists. Specifically, we examined the well-characterized CWxP, PIF, NPxxY, DRY motifs, and the sodium ion-binding site, all of which have been extensively validated through structural analyses of GPCRs.[5, 13, 14] These motifs are fundamental to GPCR receptor activation and are expected to emerge as key components in inferred allosteric paths. Notably, distinct motifs exhibit differential roles in receptor activation and inactivation: CWxP and PIF are primarily associated with receptor activation[14], whereas the NPxxY and DRY motifs, along with the sodium ion-binding site, are predominantly linked to the stabilization of the inactive state[14].

As illustrated in Figure 2, *MDPath* effectively identifies key motifs across all evaluated test systems (Fig. 3A). Motif occurrence varies depending on receptor-ligand combinations. A striking example of the established roles of CWxP and PIF in activation emerges from the β_2_-adrenoceptor systems complexed with the agonist salbutamol and the inverse agonist carazolol. This is contrasted with the NPxxY and DRY motifs, which stabilize inactive conformations and lose most of their contacts upon activation. In these systems, motif occurrences exhibit a near-perfect inverse pattern (Fig. 3A).[5, 14] For the salbutamol-bound active state of the β2-adrenoceptor, we identify recurrent allosteric paths traversing W286^6.48^ of the CWxP motif (Fig. 3B), alongside frequent paths through I121^3.40^ and F282^6.44^ of the PIF motif. Notably, I121^3.40^ appears to function as a path convergence point, integrating multiple distinct paths originating from the extracellular domain of the receptor (Fig. 3C). In the carazololbound inactive state, we observe highly correlated allosteric paths spanning the entire NPxxY (Fig. 3D) and DRY motifs (Fig. 3E). Additionally, we identify distinct paths between R131^3.50^ of the DRY motif and E268^6.30^ (Fig. 3F), highlighting a conserved interaction within the DRY motif that stabilizes inactive-state GPCRs.[5, 14, 36]

**Fig. 2.**
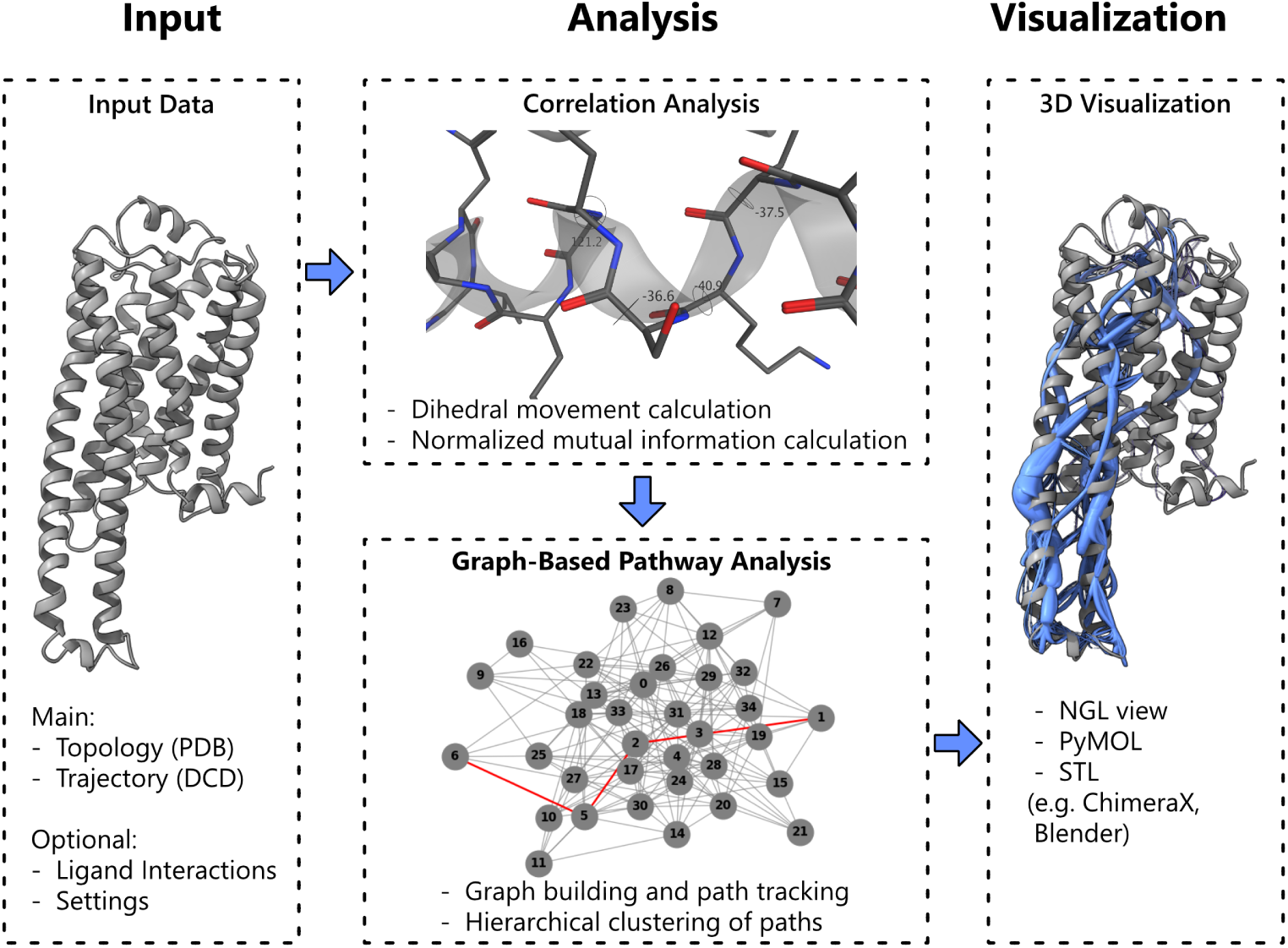
Main workflow of *MDPath* for allosteric communication path detection. The workflow starts with the input of an MD simulation, consisting of a PDB file (topology) and a DCD file (trajectory). Additional optional inputs such as interaction points for directed path tracking and settings for analysis refinement can be added. Next, *MDPath* computes residue φ-dihedral angle movements and derives normalized mutual information between residues based on them. A graph is then constructed, where residues serve as nodes, and paths are identified by maximizing normalized mutual information along the graph edges. Finally, *MDPath* generates visualization output files compatible with various molecular visualization tools, including *NGLView* [25], *PyMOL*[26], and *ChimeraX* [27].

**Fig. 3.**
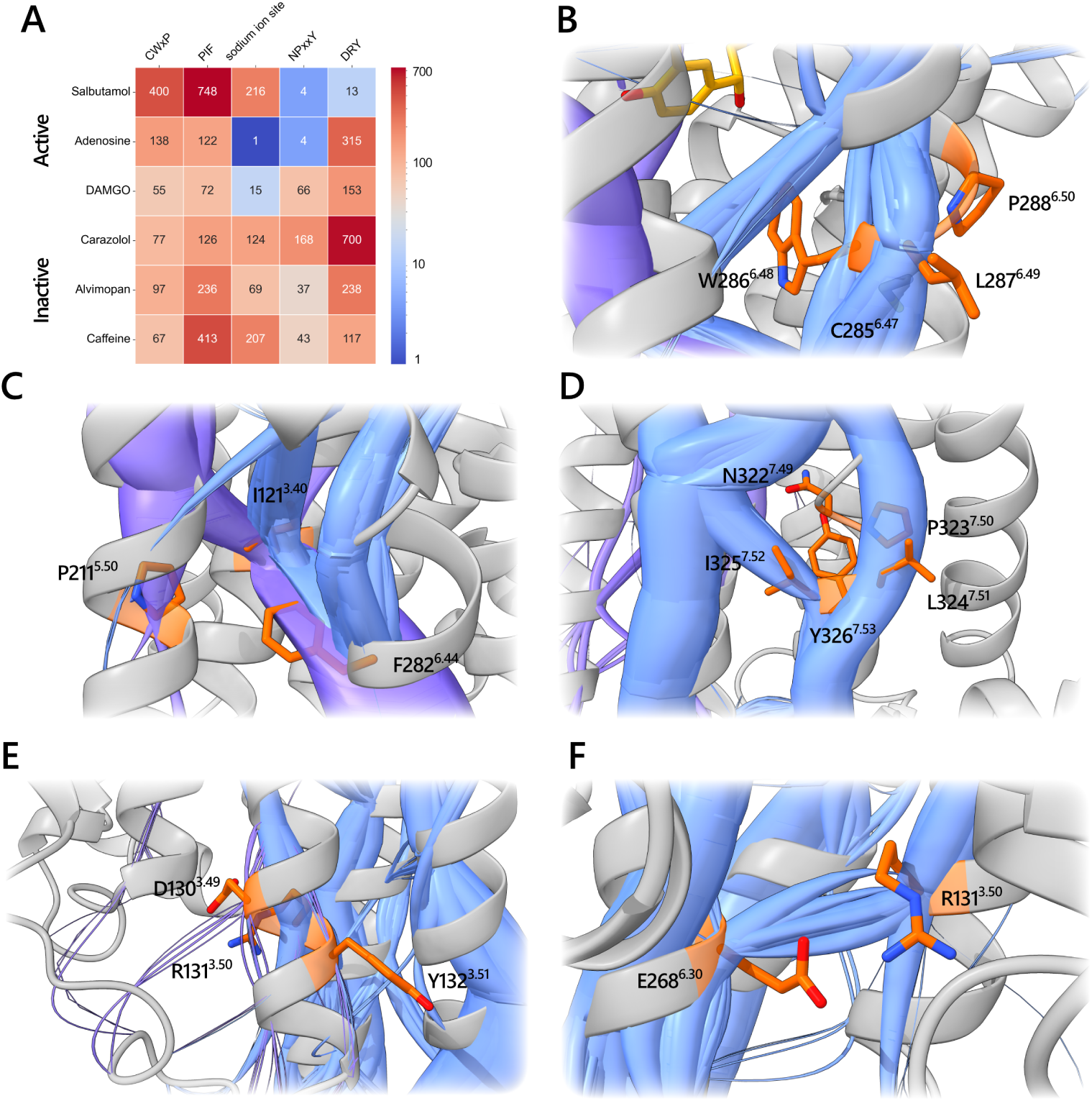
**A.** Heatmap showing involvement of conserved class A GPCR motive residues within the top 500 identified paths of all three simulation replicas. Each partition represents the specific motive’s corresponding occurrence (number) over all three simulation replicas. Heatmap color scale is plotted logarithmically to show differences more nuanced. **B.** Representation of the CWxP motive (orange) in the salbutamol bound to the β_2_-adrenoceptor. As to be expected for the agonist salbutamol strongly weighted paths (blue) are seen throughout this motive represented by the wide radius of the spline **C.** Representation of PIF motive orange) paths in the β-_2_-adrenoceptor bound to salbutamol. As for CWxP motive strongly weighted paths (blue and purple) occur through the PIF motive. Interestingly P221^5.50^ seems to be less involved in the observed activation paths, while especially I121^3.40^ acts as a path-distributing hub. **D.** Inactivation paths (blue) trancducing through the NPxxY motive (orange) in the carazolol bound β_2_-adrenoceptor. As expected for the inverse agonist bound inactive state GPCR the NPxxY motive is highly involved in path. **E.** Highly correlated paths (blue) of the D130^3.49^ and R131^3.50^ of the DRY motive (orange) in the carazolol bound β_2_-adrenoceptor, resembling the stong link of the ionic lock during the inactive state. **F.** Representation of paths (blue) between the DRY motives R131^3.50^ (orange) and E^6.30^ (orange) in the carazolol bound β_2_-adrenoceptor, displaying the second stable ionic lock formed by the DRY motive in the inactive state GPCR.

A similar situation is observed in the A_2A_-adenosine receptor bound to the endogenous agonist adenosine, which shows strong involvement of CWxP and PIF, with minimal engagement of the sodium ion site and NPxxY. The DRY motif shows enhanced traversal in this system, with dominant interhelical paths along TM3. Paths leading to TM5 or TM6 occur through the DRY motif or preceding residues. This may be due to a unique property of the A_2A_-adenosine receptor, previously noted by Hauser *et al.*[14], where the TM3 domain remains untilted during activation, facilitating this unique allosteric path. In contrast to the agonist and inverse agonist-bound structures, the A_2A_-adenosine receptor in complex with caffeine exhibits uniform envolvement of conserved motives in paths, consistent with caffeine’s role as a neutral antagonist. Visual inspection of *MDPath*-generated paths reveals a blend of patterns characteristic of both agonist and inverse agonist-bound structures, reflecting caffeine’s lack of conformational state preference. The observation of the PIF motif’s substantial involvement, formerly linked to active state paths, in simulations initiated from the inactive state, can be attributed to the comparatively high basal activity of the A_2A_-adenosine receptor. This basal activity plays a key role in the physiological tonic regulation of the central nervous system.[37]

In the DAMGO-bound MOR, we observe a distinct allosteric path pattern characterized by reduced involvement of conserved motifs compared to other systems. This can be attributed to a hub at Y328^7.43^ above the NPxxY motif being a bottleneck for paths, highlighting a unique mode of allosteric communication in the MOR. Although alvimopan is classified as an antagonist in the literature, our analysis reveals similarities to the inverse agonist bound β_2_-adrenoceptor (carazolol). This may be because alvimopan’s classification is based on its functional outcome. The lower involvement of the NPxxY motif can be attributed to the same Y328^7.43^ hub as within the DAMGO stabilized state, thus transferring the paths above the NPxxY motif lowering its occurrence within the observed paths.

To assess the statistical uniformity of the paths, 500 bootstrap samples of the dihedral angle trajectories were generated. The standard error calculated from the top 500 paths ranged from 0.89 to 2.80, indicating uniformity of the sample. This suggests that the underlying features used for path construction, correlated dihedral motions, are internally coherent within each simulation, reflecting a single conformational state rather than a mixture of multiple states. This is consistent with expectations, as no major conformational shifts were expected within the 200 ns window due to prior energy minimization.

**Fig. 4.**
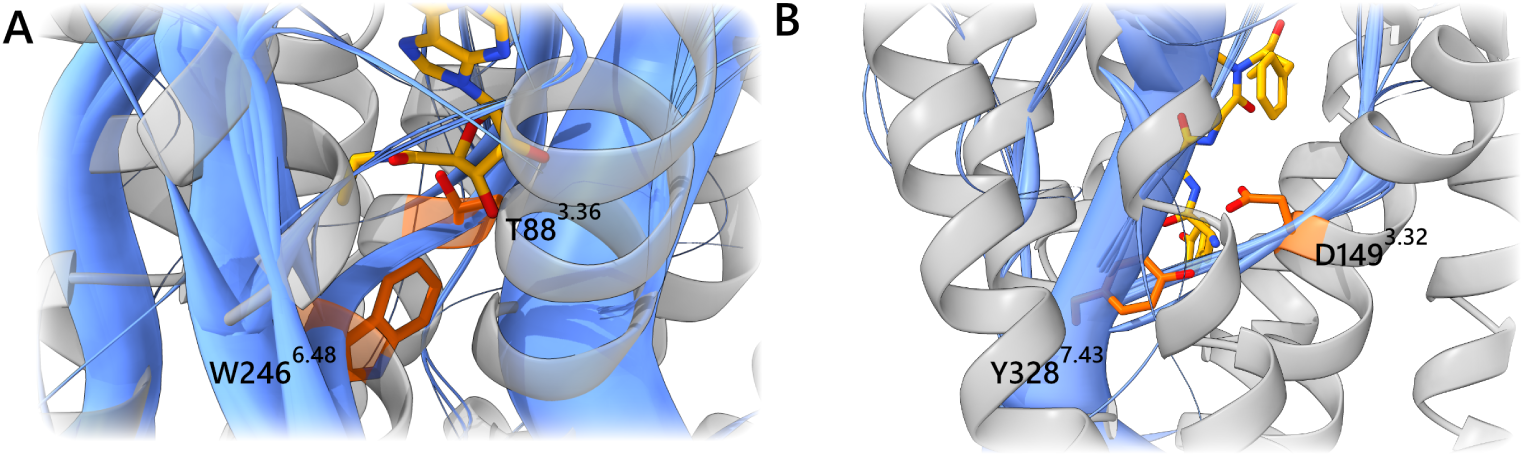
**A.** Paths (blue) going through T88^3.36^ (orange) to W246^6.48^ (orange) in the adenosine A_2A_ receptor bound to adenosine (yellow), showing the connection of residue T88^3.36^ to the known activation motive CWxP. This shows the importance of T88^3.36^ during activation allosteric paths and thus explains why mutation leads to reduced cAMP accumulation. **B.** Representation of paths (blue) through the MOR-specific path hub at position Y328^7.43^ (orange) in the MOR bound to DAMGO (yellow). This residue acts as a core distributing residue for paths towards the intracellular portion of the MOR. Thus showing the importance of this residue for MOR allosteric paths and explaining reduced activity of the MOR when mutating this residue

### 2.3 MDPath Links Receptor Mutations to Changes in Activation

To validate *MDPath*’s ability to predict receptor-specific allosteric communication paths, we investigated mutational data to assess path identification. Mutations in key residues are known to significantly impact receptor activity, offering valuable insights into activation mechanisms. We focused on mutations that affect downstream signaling, such as cAMP accumulation via the preferentially Gs-coupled receptors β_2_ and A_2A_ or cAMP reduction via the Gi-coupled MOR, because they are expected to perturb the allosteric communication paths. By comparing these mutations with paths computed by *MDPath* in the unmutated state, we assessed the accuracy of *MDPath*’s predictions in specific receptors.

For the β_2_-adrenoceptor, mutations at E268^6.30^ and D130^3.49^ that neutralize charged residues result in elevated agonist-independent cAMP levels, indicative of constitutive activity. Specifically, the D130^3.49^N, E268^6.30^Q, and E268^6.30^A mutations, as well as their combinations, induce constitutive activity, as evidenced by significantly increased cAMP accumulation in the absence of ligand stimulation.[36] In simulations of the inactive carazolol-bound state, we observe strong paths between D130^3.49^ and R131^3.50^(Fig. 3E), as well as between R131^3.50^ and E268^6.30^(Fig. 3F), suggesting that these interactions stabilize the DRY motif in the inactive state. Notably, these allosteric paths are nearly absent in the active conformation, further supporting their role in maintaining the receptor’s inactive state. The L124^3.43^R mutation, known to induce constitutive activity and elevate basal cAMP levels,[38] plays a pivotal role in the inactivation paths of the carazolol-bound β_2_-adrenoceptor. This residue serves as a critical link between two major structural clusters within the receptor, forming a path to L275^6.37^. Notably, the path between L275^6.37^ and L124^3.43^ runs parallel to the ionic lock between R131^3.50^ and E268^6.30^, suggesting a potential role in modulating receptor conformational dynamics and stabilizing the inactive state. Furthermore, mutations near the ligand-binding site of the β2-adrenoceptor, such as S203^5.42^A, S204^5.43^A, and S207^5.46^A, drastically reduce ligand binding and cAMP generation, decreasing potency by over 100-fold (S203^5.43^A: 361 nM, S204^5.44^A: 1010 nM, S207^5.46^A: 430 nM) compared to the wild-type (7.92 nM).[39] These residues, which form strong interactions with salbutamol in our simulations, are also integral to broader activation paths that converge on conserved microswitches such as the PIF motif. This elucidates how crucial ligand binding residues can also be involved in allosteric communication paths and thus directly connect binding and activation.

In a more global approach, we mapped A_2A_ receptor mutations associated with decreased activity for adenosine, taken from the GPCRdb[40], directly map to identified paths. Several mutations that affect receptor signaling within the ECL2 loop region have been characterized: mutation of C146^ECL2^ results in complete abolishment of activity measured by cAMP accumulation, whereas mutations in C159^ECL2^ and C166^ECL2^ are associated with a 21-fold and 142-fold decrease in cAMP accumulation, respectively. C146^ECL2^ is the terminal source in ECL paths identified by *MDPath*, linking to L78^3.26^ and a hub on V171^ECL2^, while C159^ECL2^ channels paths to V164^ECL2^, a major downstream path hub, and C166^ECL2^ forms a path that connects to another hub at F168^ECL2^ causing downstream paths[41]. A T88^3.36^ mutation, associated with an 83-fold decrease in cAMP accumulation and a 7.5-fold decrease in adenosine binding, directly links the TM3 path to the conserved state switch at W246^6.48^[42], providing a rational explanation for the role of T88^3.36^ in agonist-induced activation, which could not be mechanistically explained previously[42]4. As for caffeine, while mutations affect its binding to the A_2A_ receptor[43], no known mutations directly impact its activity, consistent with its role as a neutral antagonist.

Within the MOR, prior studies have linked mutations in proximity to the ICL3 at positions R278^6.31^, R279^6.32^ and R275^6.28^ with reduced inhibition of adenylate cyclase activity by DAMGO.[44] Our analysis shows that significant paths traverse towards ICL3 crossing the identified residues. In addition to the R275A^6.28^ mutation Chaipatikul *et al.* also identified I280^6.33^A to decrease the activity of DAMGO, which was also transvesed within this allosteric path through TM6.[45] A mutation at S331^7.46^A has also been identified to decrease activity during DAMGO binding.[46]S331^7.46^ demonstrates correlated movements with T120^2.54^, which establishes a connection between paths from the TM2 to TM7, subsequently influencing the NPxxY motif. As previously mentioned during the analysis of conserved motives, the path hub at position Y328^7.43^ has been shown to function as gate residue. Experimental evidence supports this, as induction of the Y328^7.43^F variant resulted in a 29-fold decrease in DAMGO binding.[47] Moreover, research suggests that mutation Y328^7.43^F not only negatively effects DAMGO binding, but also plays a role in its ability to recruit β-arrestin, which was completely lost upon mutation although G^i^-protein based signaling was still recorded.[48] A noticable feature of this residue’s presence is its consistency in the alvimopan path context, as elucidated in the analysis of conserved motifs. However, this residue contributes to a relatively minor proportion of the paths in alvimopans allosteric coupling, and the impact of mutation on MOR antagonists was comparatively less substantial.[47]

### 2.4 MDPath Enables the Mapping of Ligand-Induced Allosteric Paths

We explored MDPath’s novel capability to identify residue-specific allosteric paths, advancing beyond previous NMI-based tracking approaches[8, 10, 11]. MDPath reveals how individual residues, particularly protein-ligand contacts, influence allosteric coupling, providing mechanistic insights into the role of specific binding site amino acids. By specifying interacting residues using the ‘-lig’ flag, MDPath ensures that only allosteric paths involving these residues are considered, highlighting their role in modulating allosteric communication. Using the β_2_-adrenoceptor bound to salbutamol and carazolol, and the tertinary MOR complexed with DAMGO and G-protein, we could reveal distinct ligand-specific allosteric paths. These findings establish MDPath as a novel tool for dissecting mechanisms in a ligand-dependent context.

Distinct allosteric paths arise from salbutamol interactions, which initiate paths toward conserved activation hubs. In the dominant path, ligand interactions at TM3 residues D113^3.32^, V114^3.33^, and V117^3.36^ connect and transmit their path to the conserved micro-switch I121^3.40^ of the PIF motif, a key regulator of receptor activation[5, 14]. Additional path branches originate from S203^5.42^ and propagate through S204^5.43^, reinforcing the link to I121^3.40^, consistent with β_2_-adrenoceptor mutational data. The ligand-contact residue F290^6.52^ transduces its path directly to W286^6.48^ of the CWxP motif, which then merges with TM3 paths at T283^6.45^. Down-stream paths proceed through TM6, ultimately culminating in ICL3, whose outward shift is crucial for receptor activation[49]. Minor paths include a TM3-mediated route through R131^3.50^ of the DRY motif, linking to Y219^5.58^, a residue known to play a role in GPCR activation[14].

Similar to salbutamol, carazolol initiates major allosteric paths from V114^3.33^ and V117^3.36^, underscoring the importance of these binding site residues in β_2_-adrenoceptor allosteric coupling. However, unlike salbutamol, I121^3.40^ does not serve as a primary path hub. Instead, paths are transduced to F282^6.44^ of the PIF motif, which now acts as a major hub, consistent with its proximity to TM3 in inactive state class A GPCRs[14]. Additionally, a strong downward path through TM3 connects to the DRY motif, with high correlation between D130^3.49^ and R131^3.50^, indicative of a stabilized ionic lock characteristic of the inactive state. Another highly frequent path originates from Y316^7.43^, an interaction absent with the agonist salbutamol. This path propagates directly through TM7 to the NPxxY motif, which remains unengaged in the salbutamol-bound state, highlighting the role of this interaction in stabilizing the inactive conformation

Interestingly, in both salbutamol- and carazolol-bound β_2_-adrenoceptor analyses, we find that major allosteric paths leading to key motifs are not significantly influenced by the interaction with N312^7.39^. This suggests that while N312^7.39^ may contribute to ligand binding, its impact on functional outcomes is limited. This is further supported by the observation that both agonists and inverse agonists share similar motifs interacting with this residue, yet produce opposite functional effects. Extending this hypothesis, D113^3.32^ may also primarily serve a role in ligand binding, as it interacts with the same ligand moiety. Its involvement in allosteric paths likely arises from its proximity to V114^3.33^ and V117^3.36^, both of which prove to be essential for major paths.

DAGMO induced allosteric paths originate from the extracellular loops, where protein-ligand contacts C219^ECL2^ and W^ECL1^ act as path hubs. Paths can either take a direct route through TM3 and TM2 via C142^3.25^, I146^3.29^, or N129^2.63^, connecting to Q126^2.60^, leading to the large hub Y328^7.43^, which also represents a protein-ligand interaction, or connect directly to it via the protein-ligand interaction of D149^3.32^, which further connects the NPxxY motive with TM6. From Y328^7.43^, downstream movement at the NPxxY motif is controlled, as well as connections to the TM6 domain, with Y338^7.53^ (NPxxY) thereby interacting with the R167^3.50^ (DRY). Alternatively, another path from ECL2 connects to TM5 and TM6 via TM3, where connections are made via E231^5.36^ which connects to K305^6.58^, while V302^6.55^ and H299^6.52^ form another connection to TM3 via M153^3.36^ one or two helical turns above the CWxP, in line with the previous observation of less involvement. From there, intrahelical TM3 paths pass the DRY motif, which influences the ICL2 conformation, also directly via D166^3.49^ interacting with R181^ICL2^.

Results from combining receptor-ligand and G protein-receptor interactions were comparable to those from ligand interactions alone. However, the ligand-G protein path analysis revealed more refined allosteric communication path patterns, with reduced noise at path endings compared to the unspecific path analysis. While this additional refinement was notable, it was less pronounced than the improvement gained from incorporating protein-ligand interactions.

Statistical analysis using bootstrapping samples supports the assumption, that incorporating target residues into the analysis aids in defining paths. The inclusion of terminal path data through both receptor-ligand and receptor-protein contacts resulted in reduced standard errors, enhancing the recession of the analysis.(Table 4 SI)

### 2.5 Extending MDPath to Non-GPCR Systems: Applications in Kinase Networks

To validate the applicability of MDPaths to other protein classes, we applied MDPath to ABL1, a double-drugged kinase in the inhibited state. The analysis yielded two principal allosteric paths. The first path is initiated at the base of the myristate pocket. In the native state, this path is autoinhibited by the myristoylated N-terminal end of the SH3 domain.[50] The other one starts around the ATP-binding domain and consists of several finer paths that integrate into one larger pathway, with one of the finer paths starting at the base of the P-loop (Fig. 5A).

**Fig. 5.**
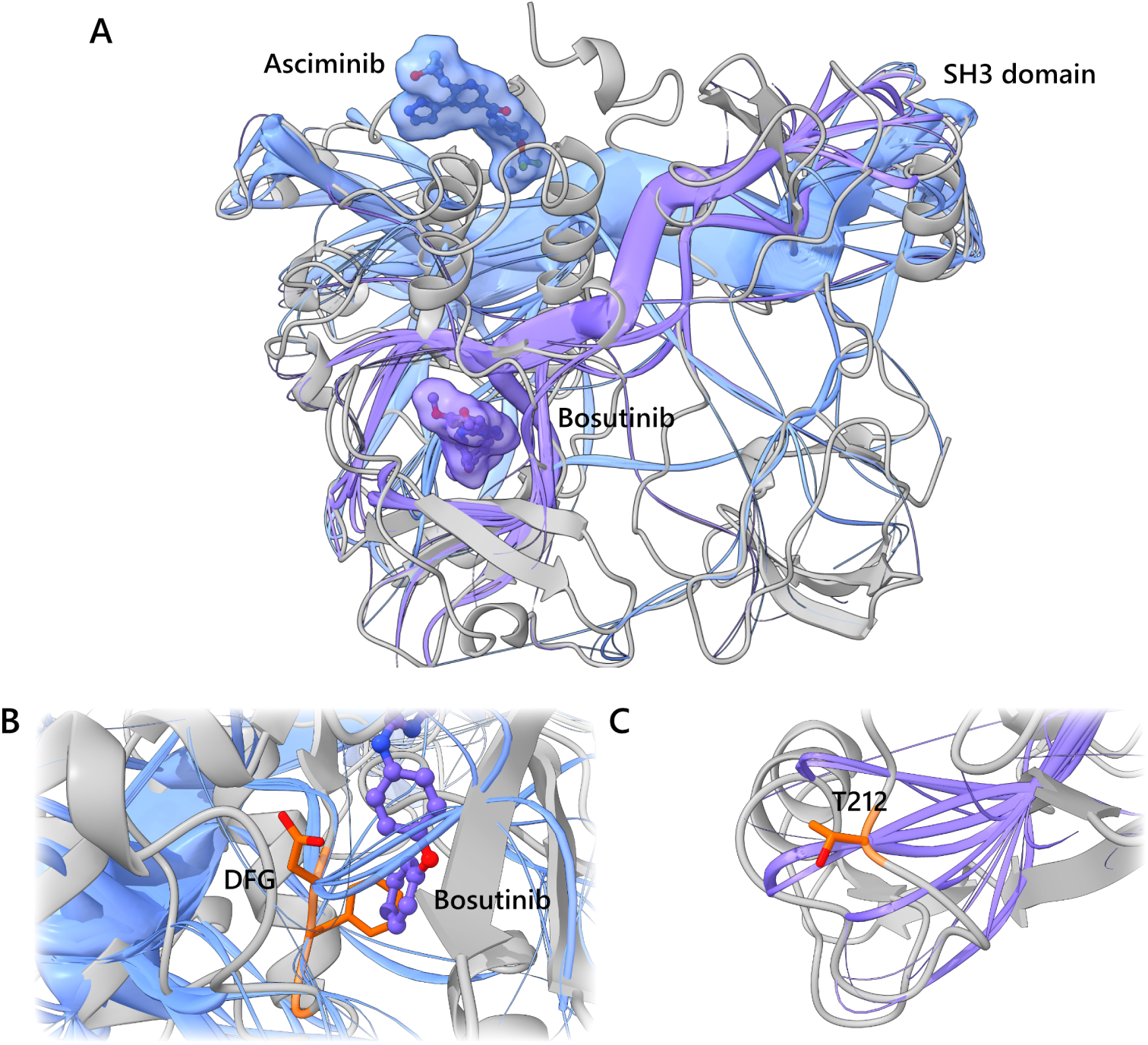
**A.** Complete view of ABL kinase bound to bosutinib (purple) and asciminib (blue). Blue paths originate from the allosteric myristate pocket bound to ascitimib and purple paths originate from the orthosteric ATP pocket bound to bosutinib. Both paths couple to the autoinhibitory SH3 domain. **B.** DFG-out motif (orange) stabilized by allosteric paths from myristate pocket (blue). This emphasizes the allosteric effect asciminib has on the orthosteric ATP-pocket in addition to effects on the autoinhibitory SH3 domain. **C.** The distant T212 (orange) is the terminal point of a path (purple) that originates from the orthosteric ATP-binding pocket, linking this residue to the ligand binding site and thus showing a correlated influence between both. Thus the molecular mechanism for reduced inhibitor activity when mutating this residue can be mechanistically explained by investigating residues along this path.

The allosteric inhibitor asciminib path transits from the myristate pocket to the ATP-binding pocket, accessing the latter’s domain via paths that affect the DFG-out motif. This provides a rational explanation for the previously noted energetic changes induced by ascitimib, which stabilized the DFG-out motif without exerting direct contact[51], thus giving a mechanistic explanation for asciminib allosteric stabilization of the inactive state at the ATP-binding pocket.

Our analysis indicates that both allosteric paths stabilize the SH3 domain via different mechanisms, demonstrating the distinct mechanisms of allosteric and orthosteric regulation of the inactive state. It is noteworthy that both paths intersect the proline-rich recognition domain, comprising residues R224, N225, K226, P227, T228, and V229 in the SH2-SH3 linker. This highlights the importance of the proline-rich recognition domain, which has been identified as a key part of SH3-induced autoinhibition in several biological systems.[52] In this particular scenario, the paths originating from the myristate pocket play a pivotal role, thereby unveiling an additional layer of the natural auto-inhibition mimicked by asciminib.

In terms of mutations, MDPath was able to identify paths that correlate with the T212R mutation far away from the known drug binding sites within the SH3 domain. Mutation of T212R has previously been shown to negatively impact clinical outcomes associated with inhibitors targeting the ATP binding pocket. Previous work found that the mutation was associated with stabilization of the active state.[53] As T212 appears to be a terminal end point in the paths starting from the orthosteric binding site of bosutinib leading to the SH3 domain. It also appears that T212 is involved in stabilizing the inactive state. Although the path appeared less distinct in the analysis due to the scaling nature of the algorithm, this could potentially underrepresent the terminal regions of the path. To address this, earlier cases in this work incorporated residue-specific data, such as protein-ligand interactions. In addition, the method could be extended to examine the effects of specific mutations on the allosteric communication paths.

The standard errors calculated for the BCR-ABL system were 1.43 on average and thus comparable to those of the GPCR systems, indicating that no major conformational changes occurred during the MD simulations. Thus, the paths within the analysis are representative of the inactive double-drugged state. (Table 4 SI)

## 3 Discussion

*MDPath* introduces a systematic workflow for analyzing allosteric communication networks, validated through comprehensive case studies of three exemplary GPCRs and the Abl kinase. The methodology not only confirms previously established activation patterns[5, 14] but also reveals ligand-specific conformational dynamics in the β_2_-adrenoceptor and MOR. By elucidating how protein-ligand interactions drive distinct structural changes, *MDPath* provides a significant advancement in understanding the molecular mechanisms of ligand-mediated protein conformational regulation.

Our validation demonstrates that the identification of conserved motifs and key residues through mutational studies was consistent with the paths identified by *MDPath*. Notably, individual replicas show nuanced differences, incorporating motifs and mutations to varying extents. This underscores the importance of generating and analyzing simulation replicas individually, as each replica may represent distinct states within the protein’s conformational ensemble.

During the investigation of the Abl kinase complexed with bosutinib and asciminib, *MDPath* successfully linked allosteric communication paths from both the orthosteric ATP pocket and the allosteric myristate pocket to distant domains. Notably, *MDPath* provides the first mechanistic insight into asciminib’s inhibitory mechanism by connecting its interaction site to the DFG motif. This exemplary analysis demonstrates the versatility of *MDPath*, proving it applicable to a wide range of protein systems. However, careful adjustment of analysis settings may be necessary for systems beyond the validated domains. In larger proteins or multi-subunit systems, modifications to path tracking and graph construction distances, and the number of evaluated paths may be required for optimal performance. To account for these scenarios, MDPath comes with modifiable input flags to enable these changes with minimal effort.

The integration of *MDPath* with complementary techniques, such as molecular docking, requires a nuanced approach to protein conformation analysis. In our validation, we consistently started from experimentally solved structures, which likely contributed to the stability and representativeness of the molecular states observed. Longer MD simulation times and careful evaluation of starting structures may help to sample such stable states, ensuring accurate and reliable analysis with *MDPath*.

Ultimately, MDPath proves to be a powerful tool for analyzing allosteric communication paths in proteins, opening new avenues for understanding ligand-induced conformational dynamics. MDPath can also analyze existing databases, such as GPCRmd, to gain new insights from existing data.[54]In the future, it may assist in identifying novel allosteric sites, thereby enabling the rational design of allosteric drugs. Furthermore, MDPath’s ability to dissect allosteric communication could prove valuable in analyzing probe dependency, predicting the effects of mutations, and elucidating complex phenomena such as functional selectivity in GPCRs.

## 4 Methods

### 4.1 System Setup

Structures of agonist-bound (β_2_: 7DHI[55]; A_2A_: 2YDO[56]; MOR: 8EFQ[57]) and antagonist-/inverse agonist-bound (β_2_: 5JQH[58]; A_2A_: 5MZP[59]; MOR: 7UL4[60]) receptors and bosutinib and asciminib bound ABL1: 8SSN[61] were retrieved from the PDB (RCSB.org)[62] and prepared using *MOE 2022.2* [63]. The proteins and their ligands (salbutamol, adenosine, DAMGO, carazolol, caffeine, alvimopan, bosutinib and asciminib) were isolated by removing additional proteins and ligands. For the MOR-DAMGO complex, the G-protein was included to assess the impact of the ternary complex on allosteric communication paths identified by *MDPath*. Missing GPCR loops were modeled using MOE’s loop modeler[63], and for the β_2_-adrenoceptor system, an additional system with truncated ICL3 was prepared. GPCRs were remutated to their native human wild-type sequences retrieved from UniProt (β_2_: P07550; A_2A_: P29274; MOR: P35372)[64]. Missing atoms were added, and atom clashes along with Ramachandran outliers were minimized using the AMBER14:ETH forcefield in *MOE* [63]. No Ramachandran outliers remained. Finally, chain endings were capped, and the structures were aligned according to the OPM database[65].

### 4.2 Molecular Dynamics Simulations

MD simulations of GPCR systems were prepared using *OpenMMDL*[66, 67], while the ABL1 system was built with CHARMM GUI[68]. GPCR systems were embedded in POPC lipid bilayer. All systems were solvated with a minimum padding of 10 Å in a cubic TIP3P water box containing 0.15 M NaCl. Simulations were performed using *OpenMM* [69] with the force fields AMBER14SB for proteins [70], Lipid21 for lipids[71], and GAFF2 for ligands[72]. The parameters chosen for ABL1 were identical to those of the GPCR systems, with the exception of using OpenFF[73] for ligand parameterization, as GAFF and GAFF2[72] were unable to set up asciminib correctly. Simulations were run on NVIDIA GeForce RTX 2080Ti, 3090 and 4090 GPUs (NVIDIA Corporation, Santa Clara). Each system was energy-minimized and equilibrated for 0.5 ns, followed by three independent 200 ns production runs under periodic boundary conditions in an NPT ensemble. Temperature (300 K) and pressure (1.0 atm) were maintained using Langevin dynamics, with a 2 fs time step. A total of 1000 frames per replica were recorded. Post-processing included trajectory alignment and centering around the protein was conducted using *OpenMMDL* [66, 67]. The centering and alignment of ABL1 was performed using VMD[74, 75], although not important for the MDPath analysis.

### 4.3 Dynamic Interaction Analysis

Protein-ligand interactions were analyzed using Dynophores (implemented in LigandScout[76])[77–82] to track the interactions of carazolol and salbutamol with the β_2_-adrenoceptor, and DAMGO with the MOR. Interaction frequencies were quantified across the molecular dynamics trajectories, with each replica analyzed independently.

### 4.4 MDPath

*MDPath* was run using standard settings with a distance cutoff of 12 Å for residues considered for clustering and path calculations, and a 5 Å cutoff for interacting residues in graph building. Conserved motif occurrence was determined by counting the occurrence of motif residues within the top 500 paths of the three replicas, with the occurrence of a motif defined as the maximum frequency of any single residue within the motif. Furthermore, a ligand interaction-based analysis was conducted for both β_2_-adrenoceptor systems and for the DAMGO/G_i_-protein stabilized MOR system. Residues for ligand-based path analysis were identified using *Dynophores*[77–82] as stated. The generation of mesh objects representing the paths as splines was accomplished through the utilization of *MDPaths* built-in spline generation function. The rendering of the figures presented in this work was performed using *ChimeraX* [27].

### 4.5 Statistical analysis

500 bootstrap samples were generated using randomly sampled dihedral angle motions of all residues. For each sample, the *MDPath* analysis was performed in the same manner as the non-bootstrap analysis. Standard errors were then calculated based on the occurrence of the top 500 paths considered in the analysis.

### 4.6 Coding and Writing

Coding assistance was provided by *ChatGPT* [OpenAI, L.L.C., San Francisco, CA], *Claude*[Anthropic PBC,San Francisco, CA] and *GitHub Copilot* [GitHub Inc., San Francisco, CA]. Grammar, spelling, punctuation and tone were edited with *Chat-GPT* [OpenAI, L.L.C., San Francisco, CA], *Claude*[Anthropic PBC, San Francisco, CA] and *DeepL*[DeepL SE, Cologne, Germany].

## Supporting information

all supplementary information for MDPath

## Acknowledgements

The authors thank Supriyo Bhattacharya and Nagarajan Vaidehi for their work in “Differences in Allosteric Communication Pipelines in the Inactive and Active States of a GPCR”[10], which served as a major inspiration for this project. Their study of allosteric communication in GPCRs provided a foundational basis that guided our approach and shaped the development of this research.

Calculations of the ABL1 system for this publication were performed on the HPC cluster PALMA II of the University of Münster, subsidized by the DFG (INST 211/667-1).

## Consent for publication

All authors have approved this version of this manuscript.

## Materials availability

PDB structures are available from the Protein Data Bank (RCSB.org)[62]: 7DHI[55], 2YDO[56], 8EFQ[57], 5JQH[58], 5MZP[59], 7UL4[60] and 8SSN[61]. MD simulation files generated and analyzed in this manuscript are availible from zenodo https://doi.org/10.5281/zenodo.15637794

## Code availability

The *MDPath* source code is publicly available at https://github.com/wolberlab/mdpath. Additionally, *MDPath* can be installed through the *Python Package Index (PyPI)*.

## Author contribution

The manuscript was written through the contributions of all authors. N.P.D. and M.T. contributed equally to this work, including method design, software development, system analysis, and presentation of results. M.B. and G.W. directed the studies.

## References

[1] Pinzi, L. & Rastelli, G. Molecular docking: Shifting paradigms in drug discovery. Int. J. Mol. Sci. 20 (2019).

[2] de Ruiter, A. & Oostenbrink, C. Advances in the calculation of binding free energies. Curr. Opin. Struct. Biol. 61, 207–212 (2020).

[3] Schaller, D., et al. Next generation 3D pharmacophore modeling. Wiley Interdiscip. Rev. Comput. Mol. Sci. 10, e1468 (2020).

[4] He, Y. et al. Nrimd, a web server for analyzing protein allosteric interactions based on molecular dynamics simulation. J. Chem. Inf. Model. 64, 7176–7183 (2024).

[5] Zhou, Q. et al. Common activation mechanism of class a gpcrs. eLife 8, e50279 (2019).

[6] Kang, P. W. et al. Calmodulin acts as a state-dependent switch to control a cardiac potassium channel opening. Science Advances 6 (2020).

[7] Westerlund, A. M., Fleetwood, O., Pérez-Conesa, S. & Delemotte, L. Network analysis reveals how lipids and other cofactors influence membrane protein allostery. The Journal of Chemical Physics 153 (2020).

[8] McClendon, C. L., Friedland, G., Mobley, D. L., Amirkhani, H. & Jacobson, M. P. Quantifying correlations between allosteric sites in thermodynamic ensembles. J. Chem. Theory Comput. 5, 2486–2502 (2009).

[9] Duncan, T. E. On the calculation of mutual information. SIAM J. Appl. Math. 19, 215–220 (1970).

[10] Bhattacharya, S. & Vaidehi, N. Differences in allosteric communication pipelines in the inactive and active states of a gpcr. Biophys. J. 107, 422–434 (2014).

[11] Dutta, S. & Shukla, D. Distinct activation mechanisms regulate subtype selectivity of cannabinoid receptors. Commun. Biol. 6, 485 (2023).

[12] Sriram, K. & Insel, P. A. G protein-coupled receptors as targets for approved drugs: How many targets and how many drugs? Mol. Pharmacol. 93, 251–258 (2018).

[13] Bock, A. & Bermudez, M. Allosteric coupling and biased agonism in g protein-coupled receptors. FEBS J. 288, 2513–2528 (2021).

[14] Hauser, A. S. et al. Gpcr activation mechanisms across classes and macro/microscales. Nat. Struct. Mol. Biol. 28, 879–888 (2021).

[15] McCorry, L. K. Physiology of the autonomic nervous system. Am. J. Pharm. Educ. 71, 78 (2007).

[16] Jacobson, K. A. & Gao, Z.-G. Adenosine receptors as therapeutic targets. Nat. Rev. Drug Discov. 5, 247–264 (2006).

[17] Stein, C. Opioid receptors. Annu. Rev. Med. 67, 433–451 (2015).

[18] Kurzrock, R. The molecular genetics of philadelphia chromosome-positive leukemias. N. Engl. J. Med. 319, 990–998 (1998).

[19] Lugo, T. G., Pendergast, A.-M., Muller, A. J. & Witte, O. N. Tyrosine kinase activity and transformation potency of bcr-abl oncogene products. Science 247, 1079–1082 (1990).

[20] Soverini, S., Mancini, M., Bavaro, L., Cavo, M. & Martinelli, G. Chronic myeloid leukemia: the paradigm of targeting oncogenic tyrosine kinase signaling and counteracting resistance for successful cancer therapy. Mol. Cancer 17, 1–15 (2018).

[21] Woessner, D. W. et al. A coiled-coil mimetic intercepts bcr-abl1 dimerization in native and kinase-mutant chronic myeloid leukemia. Leukemia 29, 1668–1675 (2015).

[22] Grafone, T. et al. A novel 4-anilino-3-quinolinecarbonitrile dual src and abl kinase inhibitor (ski-606) has in vitro activity on cml ph+blast cells resistant to imatinib. Blood 104, 1991–1991 (2004).

[23] Réa, D. & Hughes, T. P. Development of asciminib, a novel allosteric inhibitor of bcr-abl1. Crit. Rev. Oncol. Hematol. 171, 103580 (2022).

[24] Pluk, H., Dorey, K. & Superti-Furga, G. Autoinhibition of c-abl. Cell 108, 247–259 (2002).

[25] Nguyen, H., Case, D. A. & Rose, A. S. Nglview–interactive molecular graphics for jupyter notebooks. Bioinformatics 34, 1241–1242 (2017).

[26] Schrödinger, LLC. The PyMOL molecular graphics system, version 1.8 (2015).

[27] Pettersen, E. F. et al. Ucsf chimerax: Structure visualization for researchers, educators, and developers. Protein Sci. 30, 70–82 (2021).

[28] Richard J. Gowers et al. Sebastian Benthall & Scott Rostrup (eds) MDAnalysis: A Python Package for the Rapid Analysis of Molecular Dynamics Simulations. (eds Sebastian Benthall & Scott Rostrup) Proceedings of the 15th Python in Science Conference, 98 –105 (2016).

[29] Harris, C. R. et al. Array programming with NumPy. Nature 585, 357–362 (2020). URL 10.1038/s41586-020-2649-2.

[30] pandas development team, T. pandas-dev/pandas: Pandas (2020).

[31] Virtanen, P. et al. SciPy 1.0: Fundamental Algorithms for Scientific Computing in Python. Nat. Methods 17, 261–272 (2020).

[32] Pedregosa, F. et al. Scikit-learn: Machine learning in Python. J. Mach. Learn. Res. 12, 2825–2830 (2011).

[33] Aric A., Hagberg, D. A. S. & Swart, P. J. Varoquaux, G., Vaught, T. & Millman, J. (eds) Exploring network structure, dynamics, and function using networkx. (eds Varoquaux, G., Vaught, T. & Millman, J.) Proceedings of the 7th Python in Science Conference (SciPy2008), 11–15 (2008).

[34] Dijkstra, E. W. A note on two problems in connexion with graphs. Numer. Math. 1, 269–271 (1959).

[35] Foundation, B. Blender (2025). URL https://www.blender.org/. Software used for 3D modeling and animation.

[36] Ballesteros, J. A. et al. Activation of the beta2-adrenergic receptor involves disruption of an ionic lock between the cytoplasmic ends of transmembrane segments 3 and 6*. J. Biol. Chem. 276, 29171–29177 (2001).

[37] Ibrisimovic, E. et al. Constitutive activity of the a2a adenosine receptor and compartmentalised cyclic amp signalling fine-tune noradrenaline release. Purinergic Signal. 8, 677–692 (2012).

[38] Tao, Y.-X., Abell, A. N., Liu, X., Nakamura, K. & Segaloff, D. L. Constitutive activation of g protein-coupled receptors as a result of selective substitution of a conserved leucine residue in transmembrane helix iii. Mol. Endocrinol. 14, 1272–1282 (2000).

[39] Sato, T., Kobayashi, H., Nagao, T. & Kurose, H. Ser203 as well as ser204 and ser207 in fifth transmembrane domain of the human beta2-adrenoceptor contributes to agonist binding and receptor activation. Br. J. Pharmacol. 128, 272–274 (1999).

[40] Pándy-Szekeres, G., et al. Gpcrdb in 2018: adding gpcr structure models and ligands. Nucleic Acids Res. 46, D440–D446 (2017).

[41] De Filippo, E. et al. Role of extracellular cysteine residues in the adenosine a2a receptor. Purinergic Signal. 12, 313–329 (2016).

[42] Bertheleme, N., Singh, S., Dowell, S. J., Hubbard, J. & Byrne, B. Loss of constitutive activity is correlated with increased thermostability of the human adenosine ¡scp¿a2a¡/scp¿ receptor. Br. J. Pharmacol. 169, 988–998 (2013).

[43] Zhukov, A. et al. Biophysical mapping of the adenosine a2a receptor. J. Med. Chem. 54, 4312–4323 (2011).

[44] Wang, H. A conserved arginine in the distal third intracellular loop of the *µ*-opioid receptor is required for g protein activation. Journal of Neurochemistry 72, 1307–1314 (1999).

[45] Chaipatikul, V., Loh, H. H. & Law, P. Ligand-selective activation of *µ*-opioid receptor: Demonstrated with deletion and single amino acid mutations of third intracellular loop domain. J. Pharmacol. Exp. Ther. 305, 909–918 (2003).

[46] Pil, J. & Tytgat, J. Serine 329 of the *µ*-opioid receptor interacts differently with agonists. J. Pharmacol. Exp. Ther. 304, 924–930 (2003).

[47] Mansour, A. et al. Key residues defining the *µ*-opioid receptor binding pocket: A site-directed mutagenesis study. J. Neurochem. 68, 344–353 (1997).

[48] Hothersall, J. D. et al. Residues w320 and y328 within the binding site of the *µ*-opioid receptor influence opiate ligand bias. Neuropharmacology 118, 46–58 (2017).

[49] Sadler, F. et al. Autoregulation of gpcr signalling through the third intracellular loop. Nature 615, 734–741 (2023).

[50] Corbi-Verge, C. et al. Two-state dynamics of the sh3–sh2 tandem of abl kinase and the allosteric role of the n-cap. Proceedings of the National Academy of Sciences 110 (2013).

[51] Oruganti, B. et al. Allosteric enhancement of the bcr-abl1 kinase inhibition activity of nilotinib by cobinding of asciminib. J. Biol. Chem. 298, 102238 (2022).

[52] Zarrinpar, A., Bhattacharyya, R. P. & Lim, W. A. The structure and function of proline recognition domains. Sci STKE 2003 (2003).

[53] Sherbenou, D. W. et al. Bcr-abl sh3-sh2 domain mutations in chronic myeloid leukemia patients on imatinib. Blood 116, 3278–3285 (2010).

[54] Rodríguez-Espigares, I., et al. Gpcrmd uncovers the dynamics of the 3d-gpcrome. Nat. Methods 17, 777–787 (2020).

[55] Yang, F. et al. Different conformational responses of the Beta2-adrenergic receptor-Gs complex upon binding of the partial agonist salbutamol or the full agonist isoprenaline. Natl. Sci. Rev. 8, nwaa284 (2020).

[56] Lebon, G. et al. Agonist-bound adenosine a2a receptor structures reveal common features of gpcr activation. Nature 474, 521–525 (2011).

[57] Zhuang, Y. et al. Molecular recognition of morphine and fentanyl by the human mu-opioid receptor. Cell 185, 4361–4375.e19 (2022).

[58] Staus, D. P. et al. Allosteric nanobodies reveal the dynamic range and diverse mechanisms of g-protein-coupled receptor activation. Nature 535, 448–452 (2016).

[59] Cheng, R. K. et al. Structures of human a1 and a2a adenosine receptors with xanthines reveal determinants of selectivity. Structure 25, 1275–1285.e4 (2017).

[60] Robertson, M. J. et al. Structure determination of inactive-state gpcrs with a universal nanobody. Nat. Struct. Mol. Biol. 29, 1188–1195 (2022).

[61] Kim, C. et al. A biophysical framework for double-drugging kinases. Proc. Natl. Acad. Sci. U.S.A. 120 (2023).

[62] Burley, S. K. et al. Rcsb protein data bank (rcsb.org): delivery of experimentally-determined pdb structures alongside one million computed structure models of proteins from artificial intelligence/machine learning. Nucleic Acids Res. 51, D488–D508 (2022).

[63] Chemical Computing Group ULC. Molecular operating environment (MOE).

[64] Consortium, T. U. UniProt: the Universal Protein Knowledgebase in 2023. Nucleic Acids Res. 51, D523–D531 (2022).

[65] Lomize, M. A., Pogozheva, I. D., Joo, H., Mosberg, H. I. & Lomize, A. L. OPM database and PPM web server: resources for positioning of proteins in membranes. Nucleic Acids Res. 40, D370–D376 (2011).

[66] Talagayev, V. et al. Openmmdl - simplifying the complex: Building, simulating, and analyzing protein–ligand systems in openmm. J. Chem. Inf. Model. 65, 1967–1978 (2025).

[67] Talagayev, V., Chen, Y., Doering, N. P., Obendorf, L. & Wolber, G. Openmmdl (2024). URL https://github.com/wolberlab/OpenMMDL. (accessed 2024, December 12).

[68] Jo, S., Kim, T., Iyer, V. G. & Im, W. Charmm-gui: A web-based graphical user interface for charmm. J. Comput. Chem. 29, 1859–1865 (2008).

[69] Eastman, P. et al. Openmm 8: Molecular dynamics simulation with machine learning potentials. J. Phys. Chem. B 128, 109–116 (2024).

[70] Maier, J. A. et al. ff14sb: Improving the accuracy of protein side chain and backbone parameters from ff99sb. J. Chem. Theory Comput. 11, 3696–3713 (2015).

[71] Dickson, C. J., Walker, R. C. & Gould, I. R. Lipid21: Complex lipid membrane simulations with amber. J. Chem. Theory Comput. 18, 1726–1736 (2022).

[72] He, X., Man, V. H., Yang, W., Lee, T.-S. & Wang, J. A fast and high-quality charge model for the next generation general amber force field. J. Chem. Phys. 153 (2020).

[73] Boothroyd, S. et al. Development and benchmarking of open force field 2.0.0: The sage small molecule force field. J. Chem. Theory Comput. 19, 3251–3275 (2023).

[74] Humphrey, W., Dalke, A. & Schulten, K. VMD – Visual Molecular Dynamics. J. Mol. Graph. 14, 33–38 (1996).

[75] Stone, J., Gullingsrud, J., Grayson, P. & Schulten, K., Hughes, J. F. & Séquin, C. H. (eds) A system for interactive molecular dynamics simulation. (eds Hughes, J. F. & Séquin, C. H.) 2001 ACM Symposium on Interactive 3D Graphics, 191–194 (ACM SIGGRAPH, New York, 2001).

[76] Wolber, G. & Langer, T. Ligandscout: 3-d pharmacophores derived from protein-bound ligands and their use as virtual screening filters. J. Chem. Inf. Model. 45, 160–169 (2004).

[77] Sydow, D. Dynophores: Novel dynamic pharmacophores (2015).

[78] Bock, A. et al. Ligand binding ensembles determine graded agonist efficacies at a g protein-coupled receptor. J. Biol. Chem. 291, 16375–16389 (2016).

[79] Wunsch, F., Nguyen, T. N., Wolber, G. & Bermudez, M. Structural determinants of sphingosine-1-phosphate receptor selectivity. Arch. Pharm. 356, 2300387 (2023).

[80] Puls, K., Schmidhammer, H., Wolber, G. & Spetea, M. Mechanistic characterization of the pharmacological profile of hs-731, a peripherally acting opioid analgesic, at the mu-, delta-, kappa-opioid and nociceptin receptors. Molecules 27, 919 (2022).

[81] Schaller, D., et al. Next generation 3d pharmacophore modeling. Wiley Interdiscip. Rev. Comput. Mol. Sci. 10, e1468 (2020).

[82] Sydow, D. & Wolber, G. dynophores (2024). URL https://github.com/wolberlab/dynophores. (accessed 2024, October 28).

