## Supplementary material for "MDPath: Unraveling Allosteric Communication Paths of Drug Targets through Molecular Dynamics Simulations": all supplementary information for MDPath

### Supplementary Information for "MDPath: Unraveling Allosteric Communication Paths of Drug Targets through Molecular Dynamics Simulations"

Niklas Piet Doering,[†,‡] Marvin Taterra,[†,¶] Marcel Bermúdez,[*,¶] and Gerhard Wolber[*,‡]

†*equally contributed to this work*

‡*Department of Biology, Chemistry and Pharmacy, Institute of Pharmacy, Molecular Design Group, Freie Universität Berlin, Königin-Luisestr. 2+4, 14195 Berlin, Germany*

¶*Department of Theoretical Chemistry, Institute of Pharmaceutical and Medicinal Chemistry, Universität Münster, Correnstr. 48, 48149, Münster, Germany*

### S1 Dynophore Analysis

#### Salbutamol

Table 1: Summary of the most frequent receptor-ligand interactions between salbutamol and the $\beta_2$-adrenoceptor identified by *Dynophores*.[1–6] Key binding site residues and interaction types are listed, highlighting the dynamic nature of these interactions.

| **GPCR Residues** | **Interaction Types** |
|---|---|
| V114$^{3.33}$, V117$^{3.36}$<br>F290$^{6.52}$ | Hydrophobic |
| S203$^{5.42}$, S207$^{5.46}$<br>N312$^{7.39}$ | H-bond acceptors |
| D113$^{3.32}$, S203$^{5.42}$<br>S207$^{5.46}$,N312$^{7.39}$ | H-bond donors |
| D113$^{3.32}$ | Positive ionizable |

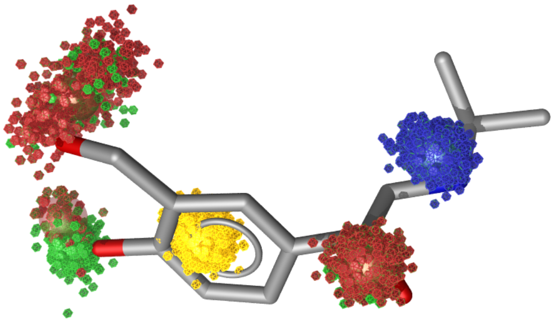

Figure 1: Point cloud representation of the salbutamol dynophore. We can see concise point clouds proving the stability of the ligand during the MD simulation. Lightly distorted clouds are seen for the phenolic hydroxy groups, as these switch their protein interaction partners between residues S203$^{5.42}$, S204$^{5.43}$ and S207$^{5.46}$. Each point within a cloud represents an interaction in one frame, with blue indicating positive ionizable interaction, red hydrogen bond acceptors, green hydrogen bond donors and yellow hydrophobic contacts.

#### Carazolol

Table 2: Summary of the most frequent receptor-ligand interactions between carazolol and the $\beta_2$-adrenoceptor identified by *Dynophores*.[1–6] Key binding site residues and interaction types are listed, highlighting the dynamic nature of these interactions.

| **GPCR Residues** | **Interaction Types** |
|---|---|
| $V114^{3.33}$, $V117^{3.36}$ | Hydrophobic |
| $F193^{ECL2}$, $T195^{ECL2}$ | |
| $A200^{5.39}$, $F290^{6.52}$ | |
| $N312^{7.39}$, $Y316^{7.43}$ | H-bond acceptors |
| $D113^{3.32}$, $S203^{5.42}$ | H-bond donors |
| $N312^{7.39}$ | |
| $D113^{3.32}$ | Positive ionizable |

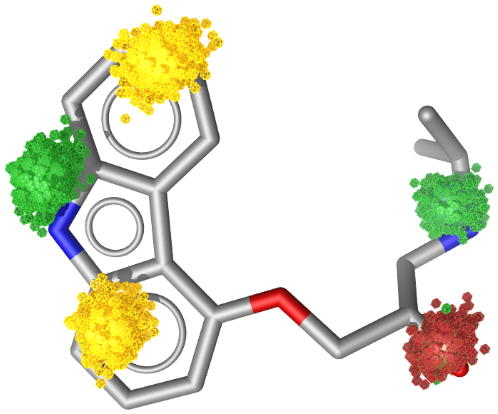

Figure 2: Point cloud representation of the carazolol dynophore. We can see concise point clouds proving the stability of the ligand during the MD simulation. Each point within a cloud represents an interaction in one frame, with blue indicating positive ionizable interaction, red hydrogen bond acceptors, green hydrogen bond donors and yellow hydrophobic contacts.

### DAMGO

Table 3: Summary of receptor-ligand and receptor-G protein interactions identified and tracked using *Dynophores*.[1–6] Key binding site residues and interaction types are listed for each ligand, highlighting the dynamic nature of these interactions.

| **Ligand** | **GPCR Residues** | **Interaction Types** |
|---|---|---|
| DAMGO | $W135^{ECL1}$, $V145^{3.28}$, $I146^{3.29}$ $Y150^{3.33}$, $M153^{3.36}$, $I298^{6.51}$ $V302^{6.55}$, $I324^{7.39}$ | Hydrophobic |
| | $Q129^{2.60}$, $H299^{6.52}$, $W320^{7.35}$ | H-bond acceptors |
| | $Q126^{2.60}$, $D149^{3.32}$, $C219^{ECL2}$ $E231^{5.35}$, $Y328^{7.43}$ | H-bond donors |
| | $D149^{3.32}$ | Positive ionizable |
| $G_i$-protein | $V171^{3.54}$, $L178^{ICL2}$, $M257^{5.61}$ $L261^{5.65}$, $V264^{5.68}$, $I280^{6.33}$ $V284^{6.37}$ | Hydrophobic |
| | $T99^{12.48}$, $R181^{ICL2}$, $R279^{6.32}$ $E343^{8.48}$, $R347^{8.52}$ | H-bond acceptors |
| | $A170^{3.53}$, $P174^{ICL2}$,$I258^{5.62}$ $R260^{5.64}$, $K262^{5.66}$ | H-bond donors |
| | $R181^{ICL2}$, $R265^{ICL3}$ | Negative ionizable |
| | $W194^{4.50}$ | Positive ionizable |

### S2 Effects of ICL3 truncation

The truncation of ICL3, as observed in several *in silico* $\beta$2-adrenoceptor studies,[7,8] produced similar signaling pathways at the binding site but struggled to replicate signals towards the intracellular regions. In our analysis of a truncated $\beta_2$-adrenoceptor in complex with salbutamol, we observe overall less defined signaling pathways. Pathways that are typically seen in activation like the connection between I3.40 to W6.48 PIW motive are still present, however the closer paths come to the truncated ICL region the more paths we see that are not linked to activation. For example we can now clearly see strong correlations of the D and R of the DRY motive while no connection to the Y is seen. This pattern would typically be indicative of inactivation and is seen in the carazolol case. During visual inspection of the truncated simulation larger movements at the truncated helix tips were observed, which may significantly affecting the NMI with residues at the intracellular side of TM5 and 6. Therefore, we conclude that meticulous modeling of the full system is crucial for accurate analysis with *MDPath*.

### S3 Bootstrapping analysis

Table 4: Standard errors within top 500 paths based on 500 bootstrap samples for each base analysis.

| **System** | **Simulation 1** | **Simulation 2** | **Simulation 3** |
|---|---|---|---|
| Active $\beta$2-adrenoceptor + salbutamol | 1.932 | 2.805 | 1.997 |
| Inactive $\beta$2-adrenoceptor + carazolol | 1.409 | 1.471 | 1.120 |
| Active A2A receptor+ adenosine | 1.471 | 2.396 | 2.617 |
| Inactive A2A receptor+ caffeine | 2.029 | 1.408 | 1.217 |
| Inactive MOR + alvimopan | 1.148 | 1.435 | 0.891 |
| Active MOR + DAMGO + $G_i$-protein | 1.636 | 1.486 | 1.307 |
| ABL1 Kinase + Ascinimib + other kinase | 1.145 | 1.776 | 1.382 |

Table 5: Standard errors within top 500 paths based on 500 bootstrap samples for each ligand-based analysis.

| **System** | **Simulation 1** | **Simulation 2** | **Simulation 3** |
|---|---|---|---|
| **Contact Based Paths** | | | |
| Active MOR + DAMGO + $G_i$-protein (DAMGO based) | 0.906 | 0.603 | 0.956 |
| Active MOR + DAMGO + $G_i$-protein ($G_i$-protein based) | 1.560 | 0.786 | 1.513 |
| Active MOR + DAMGO + $G_i$-protein (DAMGO and $G_i$-protein based) | 0.424 | 0.877 | 1.393 |
| Active $\beta$2-adrenoceptor salbutamol (salbutamol based) | 0.813 | 0.212 | 1.510 |
| Inactive $\beta$2-adrenoceptor + carazolol (carazolol based) | 0.468 | 0.075 | 0.126 |

### References


(1) Sydow, D. Dynophores: Novel Dynamic Pharmacophores. 2015; DOI: 10.18452/14267.

(2) Bock, A.; Bermudez, M.; Krebs, F.; Matera, C.; Chirinda, B.; Sydow, D.; Dallanoce, C.; Holzgrabe, U.; De Amici, M.; Lohse, M. J.; Wolber, G.; Mohr, K. Ligand Binding Ensembles Determine Graded Agonist Efficacies at a G Protein-coupled Receptor. *J. Biol. Chem.* **2016**, *291*, 16375–16389, DOI: 10.1074/jbc.M116.735431.

(3) Wunsch, F.; Nguyen, T. N.; Wolber, G.; Bermudez, M. Structural Determinants of Sphingosine-1-Phosphate Receptor Selectivity. *Arch. Pharm.* **2023**, *356*, 2300387, DOI: 10.1002/ardp.202300387.

(4) Puls, K.; Schmidhammer, H.; Wolber, G.; Spetea, M. Mechanistic Characterization of the Pharmacological Profile of HS-731, a Peripherally Acting Opioid Analgesic, at the Mu-, Delta-, Kappa-Opioid and Nociceptin Receptors. *Molecules* **2022**, *27*, 919, DOI: 10.3390/molecules27030919.

(5) Schaller, D.; Šribar, D.; Noonan, T.; Deng, L.; Nguyen, T. N.; Pach, S.; Machalz, D.; Bermudez, M.; Wolber, G. Next Generation 3D Pharmacophore Modeling. *Wiley Interdiscip. Rev. Comput. Mol. Sci.* **2020**, *10*, e1468, DOI: 10.1002/wcms.1468.

(6) Sydow, D.; Wolber, G. dynophores. 2024; `https://github.com/wolberlab/dynophores`, (accessed 2024, October 28).

(7) Neale, C.; Herce, H. D.; Pomès, R.; García, A. E. Can Specific Protein-Lipid Interactions Stabilize an Active State of the Beta 2 Adrenergic Receptor? *Biophysical Journal* **2015**, *109*, 1652–1662.

(8) Sun, X.; Ågren, H.; Tu, Y. Microsecond Molecular Dynamics Simulations Provide Insight into the Allosteric Mechanism of the Gs Protein Uncoupling from the $\beta$2 Adrenergic Receptor. *The Journal of Physical Chemistry B* **2014**, *118*, 14737–14744.
